# LLM-scCurator: Data-centric feature distillation for zero-shot cell-type annotation

**DOI:** 10.64898/2025.12.28.696778

**Authors:** Ken Furudate, Koichi Takahashi

## Abstract

Zero-shot cell-type annotation with language models degrades when marker lists are dominated by biological-noise programs (e.g., ribosomal/cell-cycle/stress). We present LLM-scCurator, a data-centric, backend-agnostic framework using pre-prompt noise masking and Gini-informed distillation to recover identity markers. Across benchmarks, LLM-scCurator improves ontology-aware hierarchical accuracy from 78.4% to 86.1% (52 clusters) over naive prompting, approaching supervised/reference-transfer performance without training data. LLM-scCurator extends to spatial transcriptomics (Visium, Xenium) for reference-free discovery of fine-grained niches.

## Main

Automated cell-type annotation in single-cell and spatial transcriptomics remains challenging, particularly for fine-grained states that are weakly separated from or absent in available references. Large language models (LLMs) offer a promising reference-free option for annotating cluster-level marker summaries, but existing approaches can be overconfident or hallucinate labels if the inputs are nonspecific^1^. Standard feature-selection outputs are often dominated by clonotype genes (TCR/Ig) and ubiquitous programs (ribosomal/mitochondrial, cell-cycle, and stress), which may obscure cell-identity signals for semantic labeling^2,3^. While recent work has focused on prompting and agentic verification^4–6^, a standardized, LLM-oriented framework to suppress recurrent biological-noise programs before prompting remains underdeveloped.

To address this gap, we developed LLM-scCurator, a data-centric framework that isolates lineage- and state-defining markers via adaptive feature distillation before prompting (**Fig. 1a**). LLM-scCurator encodes biological-noise programs as regex-based noise modules, constructs candidate marker lists from batch-aware highly variable genes (HVGs) and high-Gini markers (plus canonical sentinels), and then suppresses housekeeping, stress, and clonotype programs while rescuing rare, lineage-defining transcripts (**Fig. 1b,c**; **Extended Data Fig. 1a**; **Supplementary Table 1**). To avoid empty marker lists, LLM-scCurator incorporates fallbacks that relax filtering when no candidates remain. Users can either annotate pre-defined clusters using these marker lists (e.g., imported from Scanpy^7^ or Seurat^8^) or run a hierarchical discovery mode for coarse-to-fine labeling from major lineages to fine-grained states. LLM-scCurator operates through a backend-agnostic interface—used here with Google Gemini 2.5 Pro— compatible with cloud-hosted and local LLMs.

**Fig. 1.**
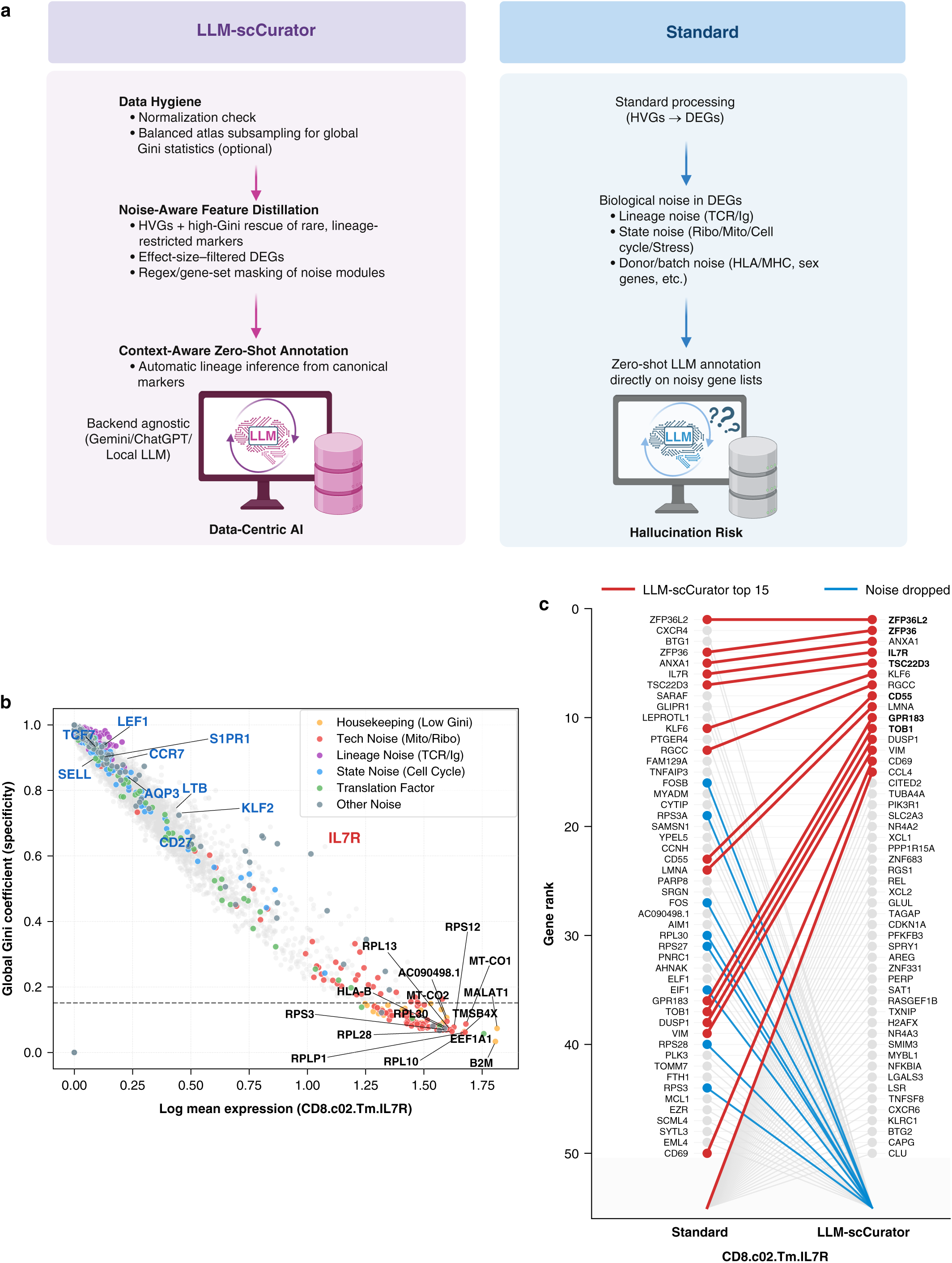
LLM-scCurator distills cluster markers before zero-shot annotation. a, Conceptual schematic comparing LLM-scCurator with a standard HVGs-to-DEGs workflow (created in BioRender). LLM-scCurator filters and rescues markers for backend-agnostic LLM prompting using regex-based noise modules with specificity-aware rescue. b, Example cluster (CD8.c02.Tm.*IL7R*): global Gini specificity versus mean expression, colored by the same keep/filter calls used to construct the prompt marker list, retaining lineage/state markers (e.g., *IL7R, TCF7,* and *CCR7*) while filtering ubiquitous programs. **Extended Data Fig. 1a** shows additional CD8⁺ clusters. c, Rank shifts between the Standard and the LLM-scCurator–distilled top 50 genes; lines connect ranks across methods and off-range genes are ghosted.

To evaluate the accuracy of LLM-scCurator for cell-type annotation, we benchmarked four tasks spanning human and mouse (CD8⁺ T cells, CD4⁺ T cells, mesenchymal differentiation [MSC], and mouse B cells). These tasks were constructed from three publicly available datasets with fine-grained annotations: a pancancer T-cell atlas^9^; an MSC subset (fibroblastic and endothelial states) from a breast cancer atlas^10^; and a nominal B-cell subset from a mouse atlas^11^, where we re-annotated mixed-lineage cells as additional lineages to test label impurity (**Extended Data Fig. 1b–i; Supplementary Tables 2 and 3**). We compared five annotation approaches—(i) LLM-scCurator; (ii) an uncurated differentially expressed genes (DEGs)-to-LLM baseline that uses the top-ranked DEGs as input without feature distillation (hereafter Standard), similar to prior GPT-based annotation^4^; and supervised/reference-transfer methods: (iii) CellTypist^12^; (iv) SingleR^13^; and (v) Azimuth^14^—and evaluated performance using an ontology-aware hierarchical accuracy metric (*S_anno_*; 0–1; **Methods**). LLM-scCurator increased mean *S_anno_* over the Standard baseline by 6.5–10.0 percentage points in the CD8⁺, CD4⁺ and MSC (no change in mouse B cells; ceiling), and matched or exceeded supervised/reference-transfer methods in the CD4⁺ and MSC benchmarks (**Fig. 2a–d; Extended Data Fig. 2a–c**).

**Fig. 2.**
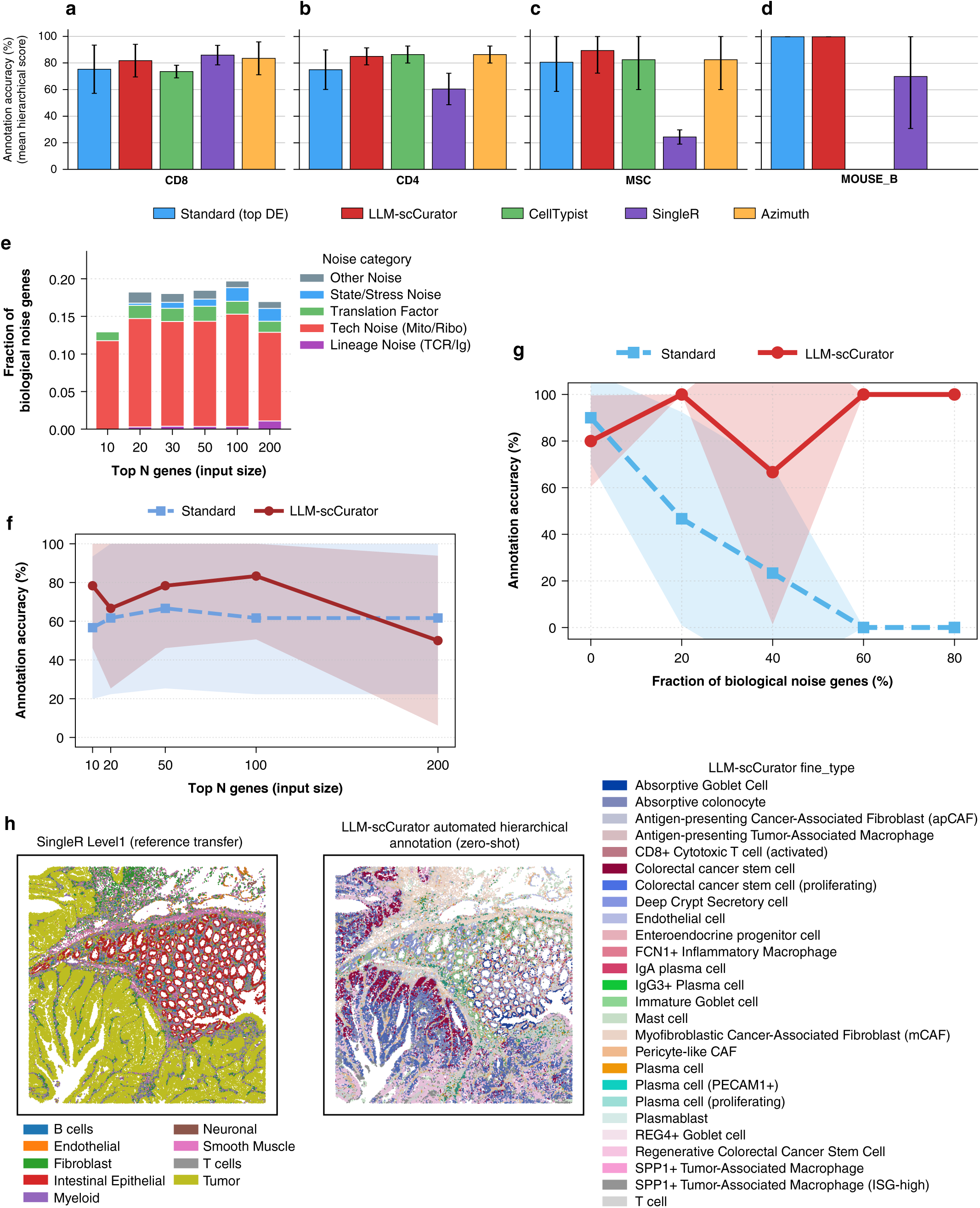
LLM-scCurator improves zero-shot annotation accuracy and robustness across modalities. a–d, Mean ontology-aware hierarchical accuracy (*S_anno_*; 95% CI across clusters) comparing Standard (uncurated DEGs-to-LLM prompting), LLM-scCurator, and supervised/reference-transfer methods across CD8⁺ (a; 17 clusters), CD4⁺ (b; 22 clusters), MSC (c; 8 clusters), and mouse B-cell (d; 5 clusters) datasets; CellTypist and Azimuth were not evaluated for mouse. e, Biological noise composition among uncurated top-*N* DEG inputs for CD8⁺ clusters (*N* = 10– 200 input genes; *n* = 17); bars show mean and 95% CI. f, Mean accuracy versus input marker-list length (N) in the CD8⁺ clusters (*n* = 6); bands indicate variability across clusters. g, Robustness to in-silico biological-noise injection in resampled 50-gene panels from CD8.c12.Tex.*CXCL13* (*n* = 3 resamplings per condition). h, Xenium colon carcinoma: SingleR Level1 label transfer (left; 9 compartments) and LLM-scCurator zero-shot coarse-to-fine labels (right; 26 epithelial/stromal states).

To assess the impact of feature distillation on LLM-based annotation, we compared LLM-scCurator with the Standard baseline by varying marker-list length and, in a separate stress test, the biological-noise fraction. Uncurated top-ranked DEGs used as the Standard inputs contained biological-noise genes (12.9–19.7% across 10–200 input genes; **Fig. 2e**). Across selected CD8⁺ clusters (*n* = 6), LLM-scCurator outperformed the Standard at 10–100 input genes (mean accuracy 66.7–83.3% vs 56.7–66.7%; **Fig. 2f**) but declined at 200 genes (50.0% vs 61.7%). With in-silico biological-noise injection in resampled 50-gene panels, LLM-scCurator remained robust (66.7–100% accuracy), whereas the Standard dropped to 0% at noise fractions ≥0.6 (**Fig. 2g**). Mechanistically, LLM-scCurator suppresses low-specificity transcripts (e.g., RPL/RPS) while preserving identity markers (e.g., *ZFP36L2* and *IL7R*; **Fig. 1b,c**), stabilizing LLM-based annotation.

Next, we asked whether LLM-scCurator enables reference-free, fine-grained spatial annotation. We applied hierarchical discovery (iterative coarse-to-fine clustering and labeling) to a Visium oral squamous cell carcinoma (OSCC) dataset^15^ and a Xenium colorectal carcinoma dataset^16^. In OSCC, LLM-scCurator subdivided pathologist-annotated regions into transcriptional programs, each supported by one-vs-rest marker effect sizes. A p-EMT/hypoxic tumor program was enriched for basal/p-EMT markers (*KRT14, KRT5, PDPN* and *LAMC2*; AUROC 0.87–1.00; log_2_FC 2.12–2.94) together with hypoxia-associated transcripts (*SLC2A1, HIF1A* and *CA9*; AUROC 0.78–0.87). A distinct basal/ISG program showed interferon activation (*STAT1* and *MX1*; AUROC 0.78–0.85). Mesenchymal/hypoxic regions were supported by extracellular matrix and mesenchymal markers (*FN1* and *VIM*; AUROC 0.85–0.86), and myofibroblastic cancer-associated fibroblasts (CAF) regions by contractile markers (*ACTA2*; AUROC 0.83; **Extended Data Fig. 3a–c**). In Xenium, SingleR Level1 label transfer from matched scRNA-seq assigned 9 compartments, whereas LLM-scCurator resolved 26 epithelial and stromal states, including stem-like (*LGR5*, *ASCL2* and *AXIN2*; AUROC 0.73–0.83; log_2_FC 1.00–2.42), deep-crypt secretory (*OLFM4* and *REG4*; AUROC 0.73–0.98), endothelial (*PECAM1* and *PLVAP*; AUROC 0.88–0.95) and pericyte-like stromal states (*RGS5* and *ACTA2*; AUROC 0.85–0.91; **Fig. 2h**; **Extended Data Fig. 3d**). Thus, LLM-scCurator enables reference-free discovery of spatially organized transcriptional programs that subdivide histology-defined regions and may capture functional differences not apparent from morphology alone.

More generally, we designed LLM-scCurator as a modular pre-prompt feature-distillation layer compatible with diverse annotation paradigms. Although recent LLM-based agents^5,6^ and a foundation-model framework^17^ differ in philosophy and implementation (**Supplementary Table 4**), they often treat marker lists as fixed inputs rather than improving them upstream. By suppressing recurrent biological-noise programs and rescuing identity markers before prompting, LLM-scCurator complements these frameworks. LLM-scCurator–distilled marker lists can serve as inputs to other LLM-based annotators and provide a less-confounded feature space for downstream analyses and reference-free spatial niche discovery.

Taken together, our results support a data-centric view of zero-shot annotation: marker-list quality is a major determinant of performance, alongside model choice. We note limitations when relevant cell types are poorly represented in an LLM’s training data and, as with other cluster-based workflows, when upstream clustering is suboptimal—especially in extremely sparse measurements. Nonetheless, LLM-scCurator improves the robustness and interpretability of zero-shot annotation by distilling less-confounded marker lists before prompting. LLM-scCurator is backend-agnostic, supports multiple LLM backends (**Supplementary Table 5**), including locally hosted models. Distributed as an open-source package with module definitions and documented filtering criteria, it provides a practical template for enforcing feature-level data hygiene in single-cell and spatial transcriptomic annotation workflows.

## Extended Data Figure legends

**Extended Data Fig. 1.**
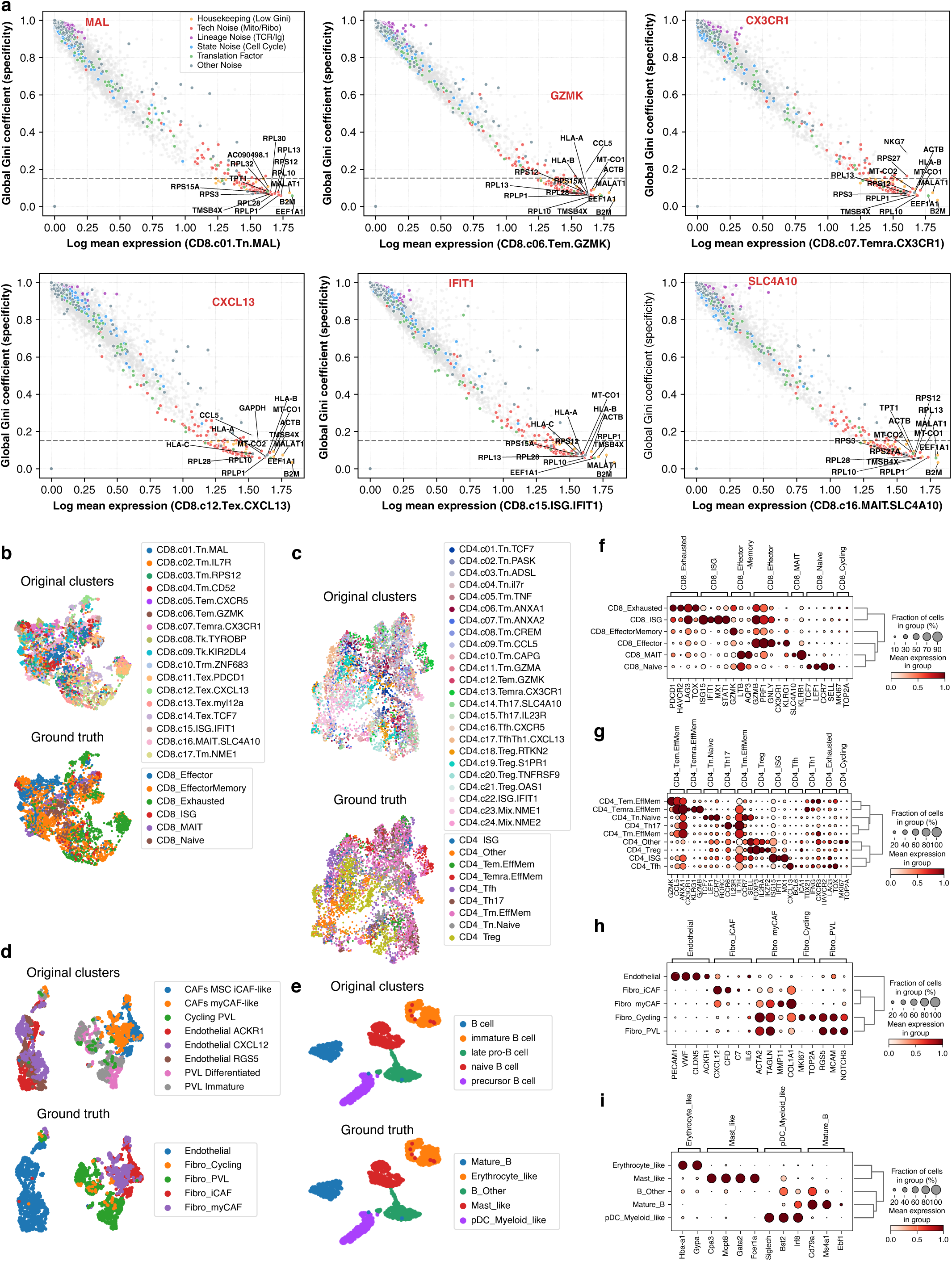
Dynamic feature masking across CD8⁺ T-cell states and harmonized ground-truth benchmarks. **a**, One panel per cluster, plotted as in **Fig. 1b** (log mean expression versus global Gini specificity) and colored by the same LLM-scCurator noise-module annotations and filtering rules used for LLM prompting. Shown clusters (as labeled in the source dataset): CD8.c01.Tn.MAL, CD8.c06.Tem.*GZMK*, CD8.c07.Temra.*CX3CR1*, CD8.c12.Tex.*CXCL13*, CD8.c15.ISG.*IFIT1*, and CD8.c16.MAIT.*SLC4A10*. **b–e,** Uniform manifold approximation and projections (UMAPs) comparing original cluster assignments (top) with the harmonized ground-truth (GT) category labels used for evaluation (bottom). Plots are shown for the stratified subsample used in benchmarking (300 cells per cluster; seed = 42). Shown datasets: pancancer CD8⁺ T cells (b), pancancer CD4⁺ T cells (c), breast cancer MSC benchmark (d), and mouse B-cell benchmark (e). **f–i,** Dot plots validating canonical marker patterns for the harmonized GT categories in each dataset (CD8⁺ [f], CD4⁺ [g], MSC [h], mouse B-cell [i]). Dot color indicates mean scaled expression and dot size indicates the fraction of cells expressing each gene. Abbreviations: Tn, naive T; Tm, memory T; Tk, cytotoxic/killer T; Th17, T helper 17; Treg, regulatory T, Mix, mixed lineage; Tem, effector memory T; Temra, effector memory RA; Tex, exhausted T; MAIT, mucosal-associated invariant T; ISG, interferon-stimulated genes; MSC, mesenchymal differentiation; CAF, cancer-associated fibroblast; iCAF, inflammatory CAF; myCAF, myofibroblastic CAF; PVL, perivascular-like.

**Extended Data Fig. 2.**
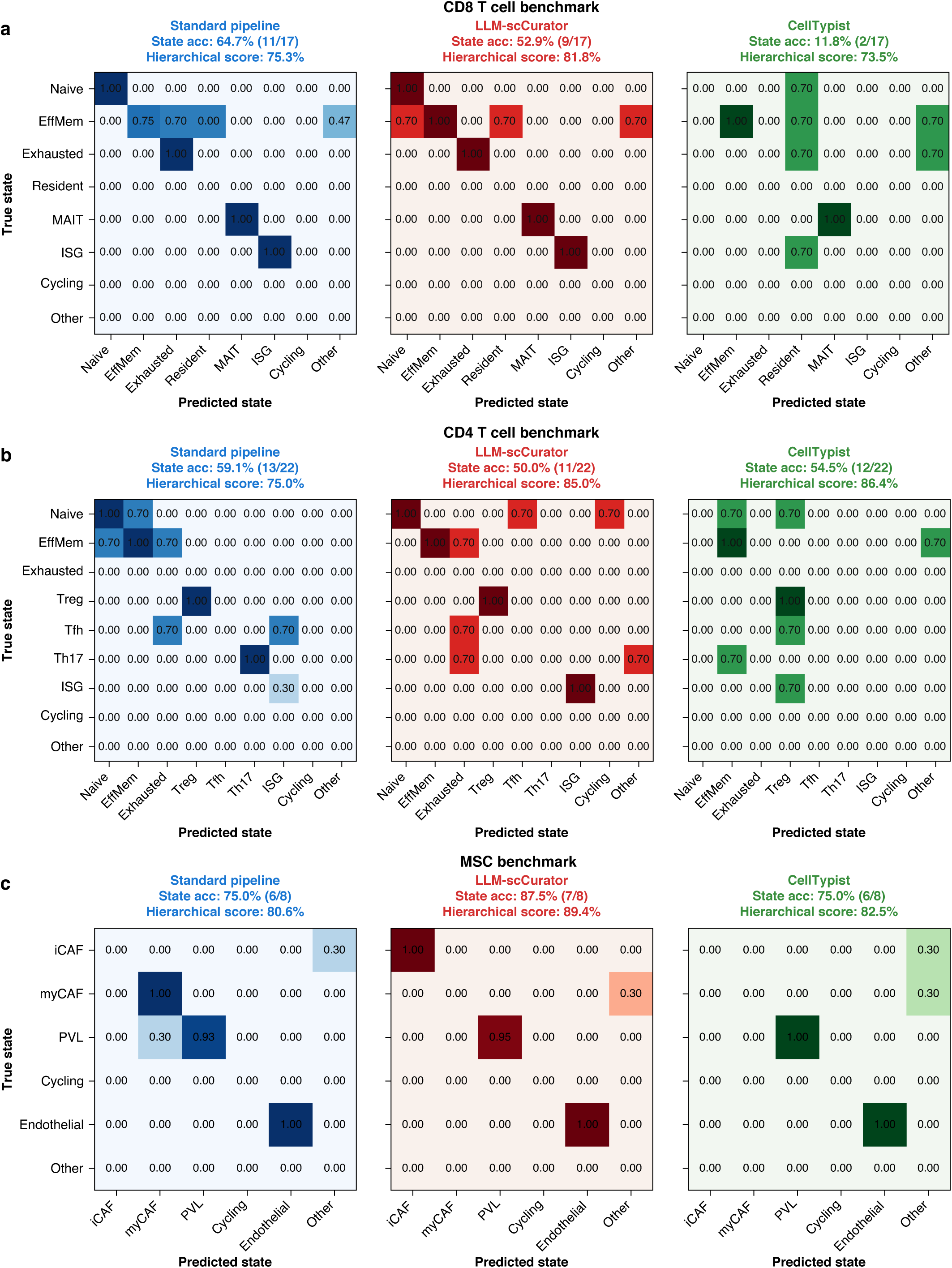
Resolution of lineage and state ambiguities by LLM-scCurator. a-c, Confusion matrices showing the mean ontology-aware hierarchical accuracy (*S_anno_*: 0–1) between GT states (rows) and predicted states (columns) for a, CD8⁺ T cells; b, CD4⁺ T cells; and c, MSC. For each lineage, panels compare the Standard pipeline (blue), LLM-scCurator (red) and CellTypist (green). The Standard pipeline frequently conflates biologically distinct but related states, whereas LLM-scCurator suppresses noise-driven confounders and concentrates high scores along the diagonal, achieving zero-shot precision comparable to or exceeding the supervised CellTypist model, particularly for rare subsets such as MAIT cells and specific stromal niches (iCAF versus myCAF). Abbreviations: ISG, interferon-stimulated genes; MSC, mesenchymal differentiation; CAF, cancer-associated fibroblast; iCAF, inflammatory CAF; myCAF, myofibroblastic CAF; PVL, perivascular-like; MAIT, mucosal-associated invariant T.

**Extended Data Fig. 3.**
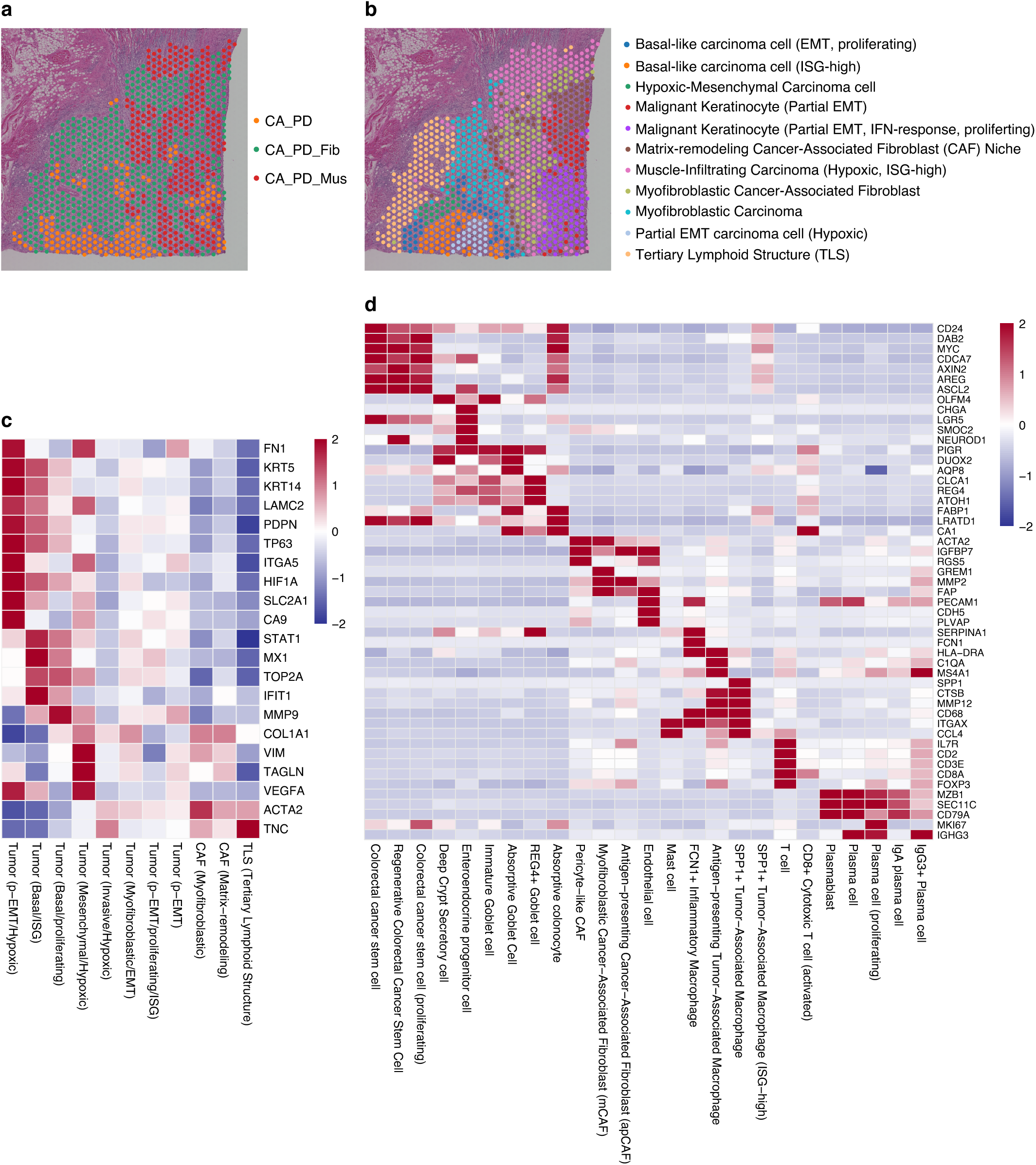
Reference-free hierarchical discovery of spatial niches in OSCC Visium and colon Xenium. a,b, Spatial maps of Visium oral squamous cell carcinoma (OSCC) comparing manual histopathology categories (a) with LLM-scCurator unsupervised hierarchical discovery (iterative coarse-to-fine clustering and labeling) on the same spots (b). Abbreviations: CA_PD, poorly differentiated carcinoma; CA_PD_Fib, CA_PD with fibrosis; CA_PD_Mus, CA_PD with muscle; CAF, cancer-associated fibroblast; EMT, epithelial–mesenchymal transition; IFN, interferon; ISG, interferon-stimulated genes; TLS, tertiary lymphoid structure. c, Pseudobulk heatmap (row-wise z-scored) across LLM-discovered OSCC niches, consistent with canonical programs (e.g., *MX1/IFIT1* in ISG-high regions, *HIF1A/VEGFA* in hypoxic niches, *PDPN/LAMC2* in p-EMT clusters). d, Xenium human colon carcinoma: pseudobulk heatmap (row-wise z-scored) across LLM-scCurator fine-type classes spanning epithelial, stromal, and immune states, consistent with known markers (e.g., *FABP1/REG4* in absorptive epithelium, *LGR5/AXIN2* in stem-like cells, *FAP/MMP2* in CAFs, *SPP1* in macrophages).

## Methods

### Study design and datasets

We evaluated four quantitative benchmarks (CD8⁺ T cells, CD4⁺ T cells, MSC and mouse B cells) derived from three publicly available atlas-style datasets with fine-grained annotations, spanning human and mouse (**Supplementary Tables 2 and 3**). Specifically, we used (i) a pancancer tumor-infiltrating T-cell atlas analyzed as separate CD8⁺ and CD4⁺ subsets across esophageal, pancreatic and renal carcinomas, (ii) a breast cancer atlas mesenchymal differentiation subset capturing fibroblastic and endothelial states, and (iii) a mouse atlas–derived nominal B-cell subset in which mixed-lineage cells were re-annotated as additional lineages (e.g., erythroid-like and mast-like), providing a controlled test for label impurity. Two additional spatial case studies (10x Genomics Visium OSCC; 10x Genomics Xenium colorectal carcinoma) were analyzed to demonstrate coarse-to-fine niche discovery (**Supplementary Table 2**).

For single-cell benchmarks, we started from author-provided count matrices and metadata and used the authors’ cluster assignments as the unit of evaluation. Ground-truth (GT) labels were standardized to harmonized categories to reduce nomenclatural variability across studies while preserving major functional axes (**Supplementary Table 2**). For the pancancer T-cell atlas, author meta-clusters were mapped to six CD8⁺ states (CD8_Naive, CD8_Effector, CD8_EffectorMemory, CD8_Exhausted, CD8_ISG, CD8_MAIT) and eight CD4⁺ states (CD4_Tn.Naive, CD4_Tm.EffMem, CD4_Tem.EffMem, CD4_Temra.EffMem, CD4_Treg, CD4_Tfh, CD4_Th17, CD4_ISG). Mappings were verified by manual inspection of canonical markers (e.g., *TCF7* for naïve states and *PDCD1* for exhausted states), and clusters lacking distinct lineage markers were excluded from benchmarking. For the breast cancer atlas mesenchymal differentiation subset, fine-grained states were collapsed into five major classes (Endothelial, Fibro_iCAF, Fibro_myCAF, Fibro_PVL, Fibro_Cycling); ambiguous clusters (Fibro_Other) were excluded from reported metrics. For the Tabula Muris Senis B-lineage subset, we re-evaluated provided annotations using lineage markers to establish a biologically consistent GT: clusters historically annotated as B cells but expressing erythroid (*Hba-a1, Gypa*) or mast-cell (*Mcpt8, Cpa3*) programs were reclassified as Erythrocyte_like and Mast_like contaminants, respectively; only clusters with canonical B-cell signatures (*Cd79a, Ms4a1*) without myeloid cross-expression were retained as Mature_B, and ambiguous B_Other clusters were excluded from benchmarking. To enable reproducible scoring of free-text LLM outputs, we defined a standardized category dictionary with GT keyword triggers and representative accepted synonyms (**Supplementary Table 4**). This dictionary was used to map model-generated labels to ontology-like standardized categories for downstream evaluation.

### Single-cell data preprocessing

All single-cell RNA-seq processing was performed in Python using Scanpy (v1.11.5). To balance population frequencies across clusters while preserving within-cluster heterogeneity, we applied stratified subsampling of *N* = 300 cells per cluster with a fixed random seed (seed = 42). Raw counts were normalized to 10,000 counts per cell and log-transformed (log[counts + 1]). Feature selection was tailored to dataset structure. For the pancancer T-cell atlas, highly variable genes (HVGs) were identified using the ‘*seurat_v3*’ flavor implementation in Scanpy with *batch_key* = *‘Cancer_Type’* to mitigate tumor-type–specific batch effects. For other single-cell benchmarks, we used the standard Seurat flavor (dispersion-based) HVG selection and retained the top 2,000 genes.

### Spatial data processing (Visium and Xenium)

Spatial workflows were adapted to the platform. For OSCC Visium, we used the processed spot-by-gene count matrix and spatial coordinates from our previously described^15^. No additional Visium-specific quality control or reprocessing was performed in this study; downstream analyses used these processed inputs as provided. For Xenium, processing was performed in R using the OSTA package (v1.2.1)^19^ (**Supplementary Table 2**). We retrieved the colorectal carcinoma dataset (Xenium_HumanColon_Oliveira) and performed quality control using scuttle. Low-quality cells were identified via adaptive thresholds based on the median absolute deviation (MAD) of library size and detected features (perCellQCFilters) and were discarded. To focus on the tumor–mucosa transition zone, we defined a region of interest (ROI) using the Xenium coordinate space (x = 2,000–5,000; y = 1,000–4,000) and subsetted cells accordingly. To account for segmentation variability, gene counts were normalized by cell area prior to downstream analyses, and principal component analysis (PCA) was used to assess global structure. The resulting processed matrix was exported to Python for LLM-scCurator analysis.

### LLM-scCurator overview and implementation

LLM-scCurator is implemented in Python and routes all model calls through a pluggable backend abstraction (*BaseLLMBackend*) exposing a single method, *generate (prompt, json_mode)*, which returns the raw text response. This isolates provider-specific SDK details from the core pipeline and standardizes how prompts and responses are handled across backends. For the analyses reported here, we used Google Gemini 2.5 Pro (*models/gemini-2.5-pro*) via GeminiBackend. Backends support an optional structured-output mode (*json_mode = True*) for steps requiring machine-readable responses; in *GeminiBackend*, this requests JSON output (*response_mime_type = ‘application/json’*) with *temperature = 0.0*. We also implemented an *OpenAIBackend* with the same interface; when supported by the API, we set *temperature = 0.0* and provided a fixed seed for increased run-to-run stability in structured-output mode.

All backend calls are wrapped with a bounded retry policy (three attempts) using exponential backoff (1, 2 and 4 seconds). When *json_mode = True*, backend exceptions are represented explicitly as a valid JSON string with an *‘Error’* label and the exception message, enabling downstream steps to record failures deterministically; these cases are treated as failed predictions in downstream evaluation.

### Feature distillation

We describe feature distillation in four parts: noise module definition, candidate-space construction from batch-aware HVGs (when batch labels are provided) and high-Gini genes (plus canonical markers) followed by DEGs and abundance/effect-size filtering, Gini-based masking with sentinel and rare-marker retention, and cross-lineage leakage control with four-stage selection and a non-empty fallback strategy. Unless stated otherwise, main analyses used default LLM-scCurator settings (top 500 DEG candidates per cluster; *N* = 50 curated markers; high-Gini candidate pool: Gini *q* = 0.9 with mean expression 0.1 relative to the global median; low-Gini housekeeping mask: Gini *q* = 0.01 capped at 0.15; cross-lineage leakage: 90th percentile-based ratio threshold with minimum 1.2 and absolute 90th percentile floor 0.1; global-context balancing: target = median group size, minimum 50 cells per group, random seed = 42).

### Noise module definitions

To reduce non-identifying biology in marker lists, we categorized confounders into lineage noise and state noise and implemented them as regular-expression (regex) patterns and curated gene lists that are compatible with both human and mouse gene nomenclature (*NOISE_PATTERNS* and *NOISE_LISTS*; **Supplementary Table 1**).

Lineage noise captures repertoire-or clonality-driven signals that can dominate DEGs without reflecting stable cell identity. We defined dedicated regex modules for T-cell receptor variable regions (human TR[ABGD][VDJ]; mouse Tr[abgd][vdj]) and immunoglobulin variable regions (human IG[HKL][VDJ]; mouse Ig[hkl][vdj]), together with immunoglobulin constant-region patterns for both species. Hemoglobin genes were also encoded as a dedicated module (Hb/Hb* patterns) to mitigate erythroid contamination signals.

State noise encompasses ubiquitous programs that frequently dominate differential expression lists, including mitochondrial genes (*MT-/mt-*), ribosomal proteins (*RPS/RPL* and *Rps/Rpl*), heat-shock genes (*HSP/Hsp*), dissociation-stress markers (*JUN/FOS* family), histone/chromatin components, and translation factors. To reduce donor-and interferon-driven confounding, we categorized MHC class I genes as state noise (human *HLA-A/B/C*…; mouse *H2-K/D/L*…), while preserving MHC class II genes for antigen-presenting cell identity. We also masked uninformative identifiers and predicted loci (ENSG/ENSMUSG, LOC, Gm, *Rik) as separate modules.

Cell-cycle programs were encoded using an ortholog-aware cycling gene list^18^ (G1/S, G2/M, and core cycling genes) and applied as a state-noise list. A small set of proliferation sentinels (e.g., *MKI67, CDK1, CCNB1/2, PCNA* and *TOP2A*; with mouse ortholog capitalization) was defined separately (*PROLIFERATION_SENTINELS*) to retain key proliferative cues when required.

### Candidate feature space, differential expression and low-expression filtering

For each cluster, we defined a candidate feature space that preserves rare, lineage-restricted markers while remaining robust to batch effects. Global gene statistics (mean log1p expression and Gini coefficients) were computed on a user-defined global reference pool set via *set_global_context*. By default, all cells in the provided atlas were used; optionally, when *balance_by* was specified, we downsampled each major group to a comparable size (target = median group size; minimum 50 cells per group when possible) to reduce dominance by abundant populations. HVGs were obtained with Scanpy (*sc.pp.highly_variable_genes*). When batch_key was provided and a raw-count layer (layers[‘counts’]) was available, we used the Seurat v3 flavor on raw counts; otherwise, we used the Seurat (dispersion-based) flavor on the current expression matrix (assumed log1p-normalized). In parallel, we augmented the candidate space with a high-Gini gene set to recover low-to-moderate abundance, but lineage-restricted markers missed by variance-based HVG selection. Genes were selected from the gene-statistics table as those in the top 10% of the Gini distribution (at or above the 90th percentile) with mean expression constrained to a bounded range (mean ≥ 0.1 and ≤ the global median; *mean_upper_quantile* = 0.5). This augmentation requires gene statistics; when a global context was not set, gene statistics were computed from the current dataset, and we recommend setting a larger atlas-level global context for benchmarking and leakage checks. The final candidate feature space for differential expression was defined as a combined set of HVGs, high-Gini genes, and a small panel of canonical lineage markers. DEGs were computed using Scanpy’s Wilcoxon rank-sum test (*sc.tl.rank_genes_groups*, “target vs rest” by default), and the top-ranked genes per cluster were used as the initial pool. We then applied explicit abundance and effect-size filters based on cluster-level mean expression and detection rates (and log fold-change when available) to suppress lowly expressed genes with negligible biological effects; if all genes were removed by these filters, we fell back to the unfiltered DEG ranking.

### Gini-based housekeeping masking and module-specific sentinel rescue

Housekeeping-like genes were defined using gene-wise global Gini coefficients computed on the global context. To avoid instability from extremely lowly expressed genes (log1p mean < 0.01), we first excluded genes below a minimum global mean-expression floor. Among the remaining genes, we masked putative housekeeping genes as those in the bottom 1st percentile of the Gini distribution (*Low_Gini_Housekeeping*; see **Fig. 1b** and **Extended Data Fig.1a**), with the effective cutoff capped at 0.15 to prevent over-masking in datasets with compressed Gini ranges. In parallel, the high-Gini candidate pool defined above was retained to ensure that rare but lineage-restricted markers remain available even when not selected by variance-based HVGs. Module-specific rescue logic was applied for LINC transcripts and hemoglobin genes. For each such module, we rank genes by global mean expression and retain only the single most highly expressed gene as a potential lineage or contamination flag, provided that its mean exceeds a minimum rescue threshold (0.1 by default); all remaining genes in the module are masked as redundant. This retains a compact sentinel for modules that may carry useful biological information (e.g., a single representative hemoglobin gene in erythrocytes or a tumor-specific LINC transcript) while preventing large, near-synonymous modules from dominating the feature space. Masking decisions are recorded as an explicit mapping from gene symbols to reason codes (e.g., *Mito_Artifact, Low_Gini_Housekeeping, Module_CellCycle_State*), enabling a transparent audit trail of excluded genes in each analysis.

### Cross-lineage specificity filtering

To distinguish lineage-specific markers from ubiquitous inflammatory or stress signatures, we implemented a data-driven cross-lineage specificity filter that operates on a balanced global context. For a given target cluster, we consider a set of candidate genes and compute, for each gene 𝑔, the 90th percentile of log1p expression within the target lineage *P*^target^_90_(*g*) and within each of the other major lineages *P*^target^_90_(*g*). We first restricted the analysis to genes with *P*^target^_90_(*g*) above a small absolute threshold, ensuring that we evaluated only genes meaningfully expressed in the target lineage. For these genes, we then examine the ratios *P*^other^_90_(*g*)/*P*^target^_90_(*g*) across all nontarget lineages and define a dynamic leakage threshold 𝑇_ratio_ as the 95th percentile of this ratio distribution, with a minimum value of 1.2.

If, in any nontarget lineage, *P*^other^_90_(*g*) < *T*_ratio_ × *P*^target^_90_(*g*), the gene is flagged as a cross-lineage contaminant (*CrossLineage_Leak*) and excluded from the final feature set. This procedure enforces a minimum degree of lineage specificity in a data-driven way and reduces the impact of markers that appear broadly in multiple lineages owing to doublets, ambient RNA or global inflammatory responses while adapting the leakage threshold to each lineage’s expression landscape.

### Four-stage feature distillation and fallback behavior

The final marker list presented to the LLM is obtained through a four-stage distillation process:

1. **Restriction to HVGs plus high-Gini and canonical lineage genes** to define a candidate space that is both batch-aware and sensitive to rare markers.
2. **Differential expression and effect-size/abundance filtering** to focus on genes whose expression differed significantly (target mean, Δmean, target detection, Δdetection, log fold change).
3. **Biological-noise program masking**, which combines regex-based modules, Gini-based housekeeping detection and cross-lineage leakage checks, yielding a set of masked genes with explicit reason codes.
4. **Mask-aware ranking and selection (with fallback),** where candidate genes are sorted by statistical significance and the list is traversed, skipping genes present in the masking map and collecting clean genes until the desired number of markers (50 by default) is reached.

In rare cases where the top-ranked DEGs are dominated by ribosomal, housekeeping or broad stress programs and all candidates are masked, LLM-scCurator reverts to progressively less stringent lists: first to the expression-filtered but unmasked DEG list and, if this is also empty, to the raw DEG ranking. This ensures that feature masking never produces a “no gene” failure for a cluster; instead, the method degrades gracefully toward a standard DEG-based input. In all cases, the masking map provides an explicit record of which genes were excluded and for what reasons. Optionally, users may provide a small gene whitelist to exempt predefined markers from masking for hypothesis-driven analyses; unless stated otherwise, all benchmarks were run without dataset-specific whitelists (beyond fixed sentinel/canonical markers) to avoid manual tuning.

### Hierarchical coarse-to-fine inference

LLM-scCurator performs annotation using a two-level hierarchical workflow. We first ensure that expression values in *adata.X* are log1p-normalized; if *highly_variable* flags are absent, HVGs are pre-computed once globally (top 2,000 by default). We then perform PCA and k-nearest neighbor graph construction and run low-resolution Leiden clustering (*coarse_res* = 0.2 and *random_state* = 42 by default) to obtain coarse clusters (*leiden_coarse*). For each coarse cluster, we curate a marker list using the full feature-distillation pipeline (including restriction of candidates to batch-aware HVGs and high-Gini genes, followed by abundance/effect-size filtering) and query the LLM in strict JSON mode without providing a parent-lineage prior. To provide biological context when a parent lineage is not specified, the implementation infers a broad lineage context from mean expression of a small canonical marker panel (e.g., *CD3* complex for T cells, *MS4A1/CD79A* for B cells, *EPCAM/KRT*s for epithelial, and *PECAM1/VWF* for endothelial) and includes it in the prompt; low-support clusters are labeled as Unknown/Mixed in this inferred context. Coarse predictions are stored as *adata.obs[‘major_type’]*, and all JSON responses (label, confidence, reasoning, and any failures) are recorded in *adata.uns[‘llm_reasoning’]* for auditability. Within each coarse cluster, we rerun PCA/neighbors and apply higher-resolution Leiden clustering (*fine_res* = 0.5 by default) to obtain fine subclusters (*leiden_fine*). Fine-level annotation reuses the same feature distillation routine, with cross-lineage leakage checks enabled using *major_type* as the coarse lineage label for global-context comparisons. The LLM is explicitly instructed to return a single JSON object of the form *{‘cell_type’: …, ‘confidence’: ‘High/Medium/Low’, ‘reasoning’: …}*, where the *cell_type* string must describe the best-matching lineage/subtype and optional state annotations are placed in parentheses (e.g., “(ISG-high)” or “(proliferating)”). JSON outputs are parsed and confidence labels are normalized to {High, Medium, Low}. Robustness to backend variability is handled by bounded retries (3 attempts with a short sleep between attempts by default). If parsing fails after all attempts, the call returns *cell_type* = ‘ParseError’ with *confidence* = ‘Low’; other repeated backend failures return *cell_type* = ‘Error’ with *confidence* = ‘Low’. In the hierarchical workflow, coarse failures fall back to *major_type* = ‘Unknown’, whereas fine-level failures fall back to the parent major label to avoid propagating empty or invalid predictions. The same Hierarchical coarse-to-fine discovery entry point is used for both single-cell and spatial datasets; for Visium spot-level data, labels are interpreted as spatial niches, whereas for Xenium cell-level data they are interpreted as per-cell types in a spatial context.

### Reproducibility and global context balancing

All analyses were performed on *AnnData* objects. LLM-scCurator includes explicit checks to distinguish log1p-normalized expression from raw counts on the basis of the dynamic range and integer likeness of *adata.X*. If the maximum value exceeds a threshold (50 by default) and the sampled entries are close to integers, the matrix is treated as raw-like. In the default mode, users are expected to supply log1p-normalized expression matrices (e.g., *sc.pp.normalize_total* followed by *sc.pp.log1p*), and the framework raises a descriptive error if a raw-like input is detected. An optional convenience mode (*allow_internal_normalization* = True) allows internal normalization: a copy of the *AnnData* object is normalized (*normalize_total* and log1p), raw counts and log-transformed values are stored in separate layers (*layers[‘counts’]* and *layers[‘logcounts’]*), and the procedure is recorded in *adata.uns[‘llm_sc_curator’][‘internal_normalization’]*. This mode is disabled by default to avoid implicit transformations of the user data. To prevent a single abundant population from dominating the global Gini distribution and cross-lineage percentiles, we compute global statistics on a balanced subsample of the atlas. Specifically, we group cells by a broad lineage label in *adata.obs[‘major_type’]* (or another user-specified column), determine the median group size, and downsample each group to this target size, with a minimum of 50 cells per group. These settings (*balance_by* = *‘major_type’*, *max_cells_per_group* = *‘auto’*, *min_cells_per_group* = 50) and the resulting numbers of input and context cells are recorded in an internal configuration object (*LLM-scCurator._global_context_config*) for reproducibility. In all the main-figure analyses, we supplied log1p-normalized matrices directly to LLM-scCurator and used the quantile-based Gini thresholds described above. For each feature distillation call, the masking module returns a per-gene map from gene ID to reason code, which we convert into tabular logs when we compute summary statistics (e.g., the fraction of TCR/Ig, ribosomal or housekeeping genes among the top-*N* DEGs in **Fig. 2e**).

### Evaluation framework and benchmarking

To quantify the impact of feature curation on LLM-based annotation, we compared an unmasked top-ranked DEGs-to-LLM prompting baseline (Standard) with LLM-scCurator. In both pipelines, we computed highly variable genes (HVGs) on the full dataset (batch-aware when batch labels were available). For each cluster, we performed a one-versus-rest differential expression test (Wilcoxon rank-sum) and ranked genes by the test statistic. In the Standard pipeline, the top-*N* ranked DEGs (𝑁 = 50 in the main analyses) were passed directly to the model. In the LLM-scCurator pipeline, we applied dynamic feature masking to ranked candidates, restricting selection to the HVG set and filtering high-Gini housekeeping genes and regex-matched lineage noise (e.g., TCR/Ig, mitochondrial and ribosomal genes). Unless otherwise noted, we disabled LLM-scCurator’s automatic context augmentation when annotating Standard DEG lists so that head-to-head differences reflect feature distillation rather than additional prompt context. We then queried Google Gemini 2.5 Pro with a fixed structured prompt to predict the predominant cell type for each cluster from the resulting gene lists. To assess robustness to input length, we repeated evaluations across list sizes (𝑁 ∈ {10, 20, 30, 100, 200}) under otherwise identical settings.

Zero-shot pipelines were benchmarked against three supervised/reference-transfer methods (CellTypist [v1.7.1], SingleR [v2.12.0] and Seurat/Azimuth [v5.4.0]). All methods used the same preprocessed data. Supervised/reference methods produced cell-level predictions that were aggregated to the cluster level by majority vote to obtain a single label per cluster. Accuracy was quantified using an ontology-aware hierarchical score (described below; **Supplementary Table 3**), where exact matches receive full credit, lineage-consistent but distinct subtypes receive partial credit, and lineage-inconsistent predictions receive zero (or low) credit.

We additionally evaluated robustness to in-silico biological-noise injection using resampled 50-gene panels from an exhausted CD8⁺ T-cell cluster (CD8.c12.Tex.*CXCL13*). We defined a noise pool by intersecting dataset genes with package-defined noise modules (regex patterns for TCR/Ig, mitochondrial and ribosomal genes; and curated gene sets including cell-cycle and other confounders). We defined a “true marker” pool using one-versus-rest DEGs for the target cluster (Wilcoxon rank-sum), excluded genes in the noise pool, and retained the top 300 candidates. For each noise fraction (0.0, 0.2, 0.4, 0.6, 0.8), we generated 50-gene panels by sampling noise genes and true markers at the corresponding proportions (three repeats; fixed random seed). Standard panels were passed directly to the LLM, whereas LLM-scCurator panels applied masking followed by replenishment with unused true markers to restore a 50-gene input. All panels used identical LLM backend and prompt settings, and predictions were scored against the cluster ground truth using the CD8 benchmark’s hierarchical scoring configuration; we report mean accuracy across repeats.

### Ontology-aware hierarchical scoring of annotation accuracy

To compare LLM-scCurator with baseline methods, we implemented an ontology-aware scoring scheme that evaluates each prediction along two axes: (i) major lineage (e.g., T cells, B cells, fibroblasts, endothelial cells, and contaminants) and (ii) functional state (e.g., naive, effector-memory, exhausted, iCAF, myCAF, and perivascular-like [PVL] states). For each benchmark, cluster identifiers were first mapped to a curated set of GT categories using deterministic rules, and the same mapping was used to derive the expected major lineage and state for every cluster. Predicted free-text labels from each pipeline (the Standard pipeline, LLM-scCurator, CellTypist, SingleR, and Azimuth) were converted into major lineages and states using dataset-specific configuration dictionaries (*HierarchyConfig* objects) that encode keyword aliases (e.g., “Temra”, “effector memory”, and “Trm” for CD8⁺ T-cell states).

Each cluster received a scalar annotation score (𝑆_anno_ ∈ [0,1]), defined as a weighted sum of lineage and state correctness,

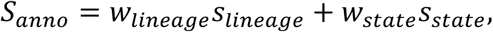

Here, 𝑠_state_ ∈ {0,1} indicates exact agreement of the predicted and expected state, and 𝑠_lineage_ ∈ {0,1} indicates agreement of the major lineage. Partial lineage credit (𝑠_lineage_ = 0.5) was enabled only for benchmark-specific, predefined near-lineage pairs (e.g., fibroblast versus endothelial in the MSC benchmark). Each benchmark also specified a set of forbidden cross-lineage predictions (e.g., predicting immune or tumor lineages for fibroblast/endothelial ground truth, or swapping T and NK lineages in the T-cell benchmarks); these predictions received a score of 0. Weights were chosen a priori to reflect the biological priority of each benchmark. For the CD8⁺ and CD4⁺ T-cell atlases, we emphasized correct T-cell lineage over within-lineage state (typically 𝑤_lineage_ = 0.7, 𝑤_state_ = 0.3) because confusing a T cell with a B cell or myeloid cell is more consequential than misclassifying closely related T-cell states. For the MSC benchmark, most clusters share a fibroblast/mesenchymal lineage and the key task is distinguishing iCAF, myCAF, PVL and cycling states; we therefore inverted the weights (e.g., 𝑤_lineage_ = 0.3, 𝑤_state_ = 0.7). For the mouse B-cell contamination benchmark, we used symmetric weights (𝑤_lineage_ = 𝑤_state_ = 0.5), as both the separation of true B-lineage cells from non-B contaminants and the correct identification of contaminant types (erythrocyte-, mast- or pDC/myeloid-like) are equally important.

The same scoring configuration was applied uniformly across methods; we report mean 𝑆_anno_ and the fraction of clusters with perfect (𝑆_anno_ = 1.0) or at least partial (𝑆_anno_ ≥ 0.5) agreement with the GT in the main figures. Evaluation code and benchmark-specific configurations are provided in the LLM-scCurator repository; canonical labels, source annotations and corresponding Cell Ontology terms are summarized in **Supplementary Table 3**.

### Spatial marker validation and effect size quantification

To summarize canonical marker support for spatial annotations, we computed per-marker effect sizes for each inferred label in a one-vs-rest manner (label versus all other spots/cells). For each gene and label, we reported (i) log_2_ fold-change (log_2_FC), log_2_[(𝜇_𝑖𝑛_ + *ε*) / (𝜇_𝑜𝑢𝑡_ + *ε*)], where 𝜇_𝑖𝑛_ and 𝜇_𝑜𝑢𝑡_ are the mean expression within and outside the label, respectively, and *ε* = 1×10^−3^. When available, log2FC was computed from a counts layer (e.g., *layers[‘counts’]*); otherwise, we used the main expression matrix. We also computed (ii) detection-rate difference (Δdet), defined as the fraction of observations with expression > *expr_threshold* (0.0 by default) in-label minus out-of-label observations; and (iii) operating characteristic curve (AUROC), which was reported as NaN for non-informative cases (zero variance or empty classes). Metrics were computed from the counts layer when available and exported as long-format CSV tables (label × gene × metrics) provided as Source Data.

### Privacy-preserving architecture and cost efficiency

LLM-scCurator is designed to minimize data sharing by performing all preprocessing and feature aggregation locally. Raw expression matrices, sample identifiers, and cell-level metadata remain in the user’s environment; only compact cluster-level summaries (ranked marker gene symbols, typically top 50, plus minimal tissue/context text) are sent to the LLM backend. This reduces exposure of individual-level measurements and decouples API usage from the number of cells; users should nevertheless follow institutional policies (e.g., IRB/DUA requirements) and the selected provider’s data-handling terms when analyzing sensitive data. Because LLM calls are issued per cluster (including coarse-to-fine steps), costs scale primarily with the number of cluster-level queries rather than dataset size. We estimated costs using a conservative budget of 1,000 total tokens per cluster query (approximately 500 input + 500 output tokens; upper bound to accommodate structured JSON responses and brief reasoning) and vendor list prices at the time of manuscript preparation (December 2025). Under these assumptions, typical runs remain low-cost across common backends, from cents for ∼50-cluster datasets to <$2 for ∼200-cluster atlas-scale analyses (**Supplementary Table 5**). Actual usage is often lower for concise JSON outputs and may vary with prompt length, model behavior, and provider billing rules.

### Data availability

All datasets analyzed in this study are publicly available from their original repositories and publications, including the pancancer T-cell atlas (GSE156728), the breast cancer single-cell atlas (GSE176078), the Tabula Muris Senis mouse atlas (figshare DOI: 10.6084/m9.figshare.8273102), and the Xenium colorectal carcinoma dataset distributed via the OSTA.data R package (*Xenium_HumanColon_Oliveira*). Processed spatial transcriptome data for OSCC Visium validation are available through the Genomic Expression Archive (GEA) under accession E-GEAD-511, which provides a processed integrated *AnnData* object *(.h5ad*) with annotations and clustering results. Supplementary spatial metadata required for selected spatial visualizations (*AnnData.uns[‘spatial’]* components) are available on Figshare (DOI: 10.6084/m9.figshare.20408067). To ensure reproducibility without redistributing primary data, we provide scripts to download and preprocess each dataset and record fixed cell/spot ID lists for benchmarking subsamples.

### Code availability

LLM-scCurator source code, benchmarking scripts, and configuration files, as well as source data underlying the figures, are available on GitHub (https://github.com/kenflab/LLM-scCurator; release tag v0.1.0) and archived on Zenodo (DOI: 10.5281/zenodo.17970494). Documentation is provided in the GitHub repository and on Read the Docs (https://llm-sccurator.readthedocs.io).

## Author contributions

K.F. conceived the study. K.F. curated the data; performed formal analysis, validation and investigation; developed the methodology; generated visualizations; and wrote the original draft. K.T. supervised the study and administered the project. K.F. and K.T. reviewed and edited the manuscript.

## References

1. Hu, M. et al. Evaluation of large language models for discovery of gene set function. Nat. Methods 22, 82–91 (2025).

2. Townes, F. W., Hicks, S. C., Aryee, M. J. & Irizarry, R. A. Feature selection and dimension reduction for single-cell RNA-Seq based on a multinomial model. Genome Biol. 20, 295 (2019).

3. Kotliar, D. et al. Identifying gene expression programs of cell-type identity and cellular activity with single-cell RNA-Seq. Elife 8, (2019).

4. Hou, W. & Ji, Z. Assessing GPT-4 for cell type annotation in single-cell RNA-seq analysis. Nat. Methods 21, 1462–1465 (2024).

5. Chen, J., Zhang, J., Yao, H. & Li, Y. CellTypeAgent: Trustworthy cell type annotation with Large Language Models. *arXiv:2505.08844*, (2025).

6. Xie, E. et al. CASSIA: a multi-agent large language model for automated and interpretable cell annotation. Nat. Commun. (2025) doi:10.1038/s41467-025-67084-x.

7. Wolf, F. A., Angerer, P. & Theis, F. J. SCANPY: large-scale single-cell gene expression data analysis. Genome Biol. 19, 15 (2018).

8. Hao, Y. et al. Dictionary learning for integrative, multimodal and scalable single-cell analysis. Nat Biotechnol 42, 293–304 (2024).

9. Zheng, L. et al. Pancancer single-cell landscape of tumor-infiltrating T cells. Science 374, abe6474 (2021).

10. Wu, S. Z. et al. A single-cell and spatially resolved atlas of human breast cancers. Nat. Genet. 53, 1334–1347 (2021).

11. Tabula Muris Consortium. A single-cell transcriptomic atlas characterizes ageing tissues in the mouse. Nature 583, 590–595 (2020).

12. Xu, C. et al. Automatic cell-type harmonization and integration across Human Cell Atlas datasets. Cell 186, 5876–5891.e20 (2023).

13. Aran, D. et al. Reference-based analysis of lung single-cell sequencing reveals a transitional profibrotic macrophage. Nat. Immunol. 20, 163–172 (2019).

14. Hao, Y. et al. Integrated analysis of multimodal single-cell data. Cell 184, 3573–3587.e29 (2021).

15. Furudate, K. et al. Spatial colocalization and molecular crosstalk of myofibroblastic CAFs and tumor cells shape lymph node metastasis in oral squamous cell carcinoma. PLoS Genet. 21, e1011791 (2025).

16. Oliveira, M. F. de et al. High-definition spatial transcriptomic profiling of immune cell populations in colorectal cancer. Nat. Genet. 57, 1512–1523 (2025).

17. Cui, H. et al. scGPT: toward building a foundation model for single-cell multi-omics using generative AI. Nat. Methods 21, 1470–1480 (2024).

## Methods References

18. Tirosh, I., et al. Dissecting the multicellular ecosystem of metastatic melanoma by single-cell RNA-seq. Science 352, 189–196 (2016).

19. Crowell, H. L., et al. Orchestrating spatial transcriptomics analysis with bioconductor. *bioRxiv*, (2025). doi: 10.1101/2025.11.20.688607.

