## Supplementary Tables for "LLM-scCurator: Data-centric feature distillation for zero-shot cell-type annotation"

**Supplementary Table 1** lists the noise modules and masking criteria implemented in LLM-scCurator—including lineage- and state-level gene sets, their sources, and activation rules—with full gene lists provided in the accompanying GitHub repository. **Supplementary Table 2** summarizes all datasets used in this study, including sample size, modality, platform, and role in each analysis. **Supplementary Table 3** details the ontology mappings and harmonization rules used to normalize author-provided labels and free-text LLM outputs into a consistent hierarchical annotation space. **Supplementary Table 4** provides a conceptual comparison of LLM-scCurator with other LLM-based annotators and single-cell foundation models, highlighting that our framework focuses on noise-aware feature distillation and is complementary to the downstream choice of the LLM backend or representation model. Finally, **Supplementary Table 5** presents a cost analysis of cluster-level annotation across varying model tiers and vendors (Google, OpenAI).

**Supplementary Table 1.** Noise modules, gene sets and masking criteria

| Noise module | Detection strategy (Source/Regex) | Masking policy & dynamic rescue |
|---|---|---|
| Lineage confounders | | |
| Adaptive immune clones | TCR/Ig variable regions (e.g., TRBV*, TRAV*, IGHV*; regex TR[ABGD][VDJ], IG[HKL][VDJ]) | Full masking. Removes V(D)J-driven clonal diversity so that the LLM does not hallucinate subtypes from specific rearrangements. Lineage itself is defined by constant regions and CD3 complex genes (e.g., *CD3E, TRAC*) retained via the lineage marker list. |
| Ig constant redundancy | Ig heavy/light constant regions (e.g., *IGHM, IGHG1–4, IGKC, IGLC1–7* and mouse orthologs) | Full masking. Prevents Ig constant transcripts from dominating plasma/B-cell profiles while plasma identity is preserved via upstream markers such as *JCHAIN, MZB1, SDC1* and *TNFRSF17*. |
| Cross-Lineage leak | Percentile-based leakage checks across major lineages (e.g., 90th-percentile expression ratios in target vs. other lineages) | Dynamic masking. Removes candidate markers when their high-percentile expression is higher in contrasting global lineages than in the target cluster (for example, Alb-like leakage in nonhepatocyte clusters). |
| Biological state | | |
| Cell cycle | S/G2M phase genes from Tirosh et al[1]. (e.g., G1/S + G2/M + core cycling set) | Selective masking. Masks bulk cell-cycle genes to unmask underlying identity while retaining key "sentinels" (e.g., *MKI67, CDK1, CCNB1/2, PCNA, TOP2A,* and *BIRC5*) so that proliferative states can still be encoded as annotation suffixes. |
| Stress & Artifacts | Mitochondrial and global stress programs (MT-, RPL/RPS, HSP*, JUN/FOS, translation factors EEF*/EIF*, histones) | Full masking. Removes signatures related to cell quality, dissociation stress and ribosomal/translation dominance so that the LLM focuses on lineage-defining markers rather than global state programs. |
| Erythrocyte contam | Hemoglobin and erythroid markers (e.g., *HBA, HBB, GYPA, ALAS2* and mouse orthologs) | Dynamic sentinel. Masks hemoglobin-driven contamination in nonerythroid clusters but retains a single top-expressed Hb gene when the mean expression > 0.1, preserving recognizability of true erythrocyte clusters. |
| Global/Technical | | |
| Global specificity | Global Gini coefficient on log1p expression (low-Gini tail among genes | Dynamic masking. Removes ubiquitous housekeeping genes with low specificity. Exceptions: genes in the |

| | | |
|---|---|---|
| | with mean ≥ 0.01; default quantile q = 0.01, capped at 0.15) | LINEAGE_MARKERS whitelist, proliferation sentinels and user whitelists are protected regardless of Gini score. |
| Technical noise | Noninformative identifiers (Ensembl IDs, LOC####, Gm####, *Rik, LINC####) | Mixed policy. Ensembl/LOC/Gm/Rik genes are fully masked as noninformative for LLM reasoning. LINC RNAs use a dynamic sentinel rule: only the single top-expressed LINC is retained when sufficiently abundant as a potential tumor- or tissue-specific marker. |
| Batch drivers | Sex- and donor-associated genes (e.g., *XIST, UTY, DDX3Y, HLA-A/B/C* and mouse *H2-D/K/L*) | Full masking. Removes sex-linked and MHC class I genes that mainly encode patient or batch effects. Human HLA class II (HLA-D*) and mouse class II (H2-A/E) are retained to preserve antigen-presenting cell definition. |

**Abbreviations**: **TCR**, T-cell receptor; **Ig**, immunoglobulin; V(D)J, variable (diversity) joining gene segments; regex, regular expression pattern; **S/G2M**, S phase / G2–M phase of the cell cycle; **MHC**, major histocompatibility complex; **HLA**, human leukocyte antigen; **H2**, mouse MHC locus; **Gini**, Gini coefficient (global gene specificity metric); **log1p**, log(1 + x) transform. **Mean expr.**, mean expression on the log1p scale. **q**, quantile threshold.

**Gene-name prefixes**: **MT-**, mitochondrial genes; **RPL/RPS**, ribosomal protein large/small subunits; **HSP**, heat shock proteins; **EEF/EIF**, translation elongation/initiation factors; **LINC**, long intergenic non-coding RNA.

**Gene-symbol conventions**: **LOC**####, predicted locus identifiers; **Gm**####, predicted mouse genes; **Rik**, RIKEN cDNA clones; **Ensembl IDs**, Ensembl gene identifiers.

**Supplementary Table 2.** Datasets and harmonized ground-truth annotations

| Dataset source | Biological context | Lineage focus | Harmonized ground-truth categories (Labels) |
|---|---|---|---|
| Pancancer T-cell atlas[2], CD8+ subset | TIL from esophageal, pancreatic and renal cell carcinomas (10x). | CD8+ T cells | Six states: CD8_Naive, CD8_Effector, CD8_EffectorMemory, CD8_Exhausted, CD8_ISG, CD8_MAIT. |
| Pancancer T-cell atlas[2], CD4+ subset | TIL from esophageal, pancreatic and renal cell carcinomas (10x). | CD4+T cells | Eight states: CD4_Tn.Naive, CD4_Tm.EffMem, CD4_Tem.EffMem, CD4_Temra.EffMem, CD4_Treg, CD4_Tfh, CD4_Th17, CD4_ISG. |
| Breast cancer atlas[3], mesenchymal differentiation subset | Stromal and vascular compartments (fibroblastic and endothelial states; hereafter, MSC) from primary breast cancers (10x) | CAF/PVL/endothelial cells | Five major classes used for benchmarking: Endothelial, Fibro_iCAF, Fibro_myCAF, Fibro_PVL, Fibro_Cycling (ambiguous Fibro_Other excluded from metrics). |
| Tabula Muris Senis[4] , B-cell subset[4] | Multiorgan mouse atlas; subset of B-cell lineage and contaminating populations. | B cells and contaminating lineages | Four classes: Mature_B, Erythrocyte_like, Mast_like, pDC_Myeloid_like (ambiguous B_Other excluded from metrics). |
| Human OSCC[5] | OSCC resection specimen profiled by Visium (10x). | Malignant and stromal niches | Pathologist-annotated tumor regions: CA_PD, CA_PD_Fib, and CA_PD_Mus. LLM-scCurator spatial labels were validated by canonical marker support (effect sizes/AUROC; Source Data) |
| Human colon carcinoma[6,7] | Colorectal carcinoma section profiled with Xenium (10x). | Epithelial, stromal and immune compartments | No GT labels. LLM-scCurator spatial labels were validated by canonical marker support (effect sizes/AUROC; Source Data) and compared to SingleR reference transfer. |

**Notes:**
For each dataset, we summarize the biological context, lineage focus and harmonized ground-truth (GT) categories used for benchmarking. GT labels were derived from author-provided annotations using deterministic string-based rules and validated with

canonical marker panels (see Extended Data Fig. 1b-i). Ambiguous “Other” categories (Fibro_Other, B_Other) were excluded from the quantitative metrics.

**Abbreviations: GT**, ground truth; **TIL**, tumor-infiltrating lymphocyte(s); **CAF**, cancer-associated fibroblast(s); **iCAF**, inflammatory CAF; **myCAF**, myofibroblastic CAF **PVL**, perivascular-like (fibroblast) cells; **pDC**, plasmacytoid dendritic cell(s); **OSCC**, oral squamous cell carcinoma; **CA_PD**, poorly differentiated carcinoma; **CA_PD_Fib**, poorly differentiated carcinoma with fibrosis; and **CA_PD_Mus**, poorly differentiated carcinoma with muscle; **10x**, 10x Genomics; **Visium**, Visium spatial gene expression; **Xenium**, Xenium in situ transcriptomics.

**Supplementary Table 3.** Label harmonization and ontology mapping

| Standardized category (ontology) | GT input keyword triggers[*] | Representative LLM output synonyms |
|---|---|---|
| Pancancer CD8$^+$ T cells (Lineage w = 0.7, State w = 0.3) | | |
| CD8_Naive | “naive”, “tn.” | naive cd8, tn, naive cytotoxic |
| CD8_EffectorMemory | “em”, “tm”, “memory”, “temra” | effector memory, tem, temra, cytotoxic effector, activated |
| CD8_Exhausted | “tex”, “exhausted”, “pdcd1” | exhausted, tex, dysfunctional, pd-1* |
| CD8_ISG | “isg”, “interferon” | interferon-stimulated, isg-high, type i ifn–responsive |
| CD8_MAIT | “mait” | mait cell, mucosal-associated invariant |
| CD8_Cycling | “cycling”, “proliferating”, “mki67”, “top2a” | proliferating, cycling, ki-67_, dividing |
| Pancancer CD4$^+$ T cells (Lineage w = 0.7, State w = 0.3) | | |
| CD4_Tn.Naive | “naive”, “tn.” | naive cd4, helper tn |
| CD4_EffectorMemory | “tem.”, “tm.”, “temra” | effector memory, central memory, tcm, tem, temra |
| CD4_Treg | “treg”, “foxp3”, “regulatory” | treg, regulatory t, foxp3* |
| CD4_Tfh | “tfh”, “follicular” | tfh, t follicular helper |
| CD4_Th17 | “th17”, “il17” | th17, il-17_ helper, rorc* |
| CD4_ISG | “isg”, “interferon” | interferon-stimulated, isg-high |
| Breast cancer CAF/MSC[†] (Lineage w = 0.3, State w = 0.7) | | |
| Fibro_iCAF | “icaf”, “msc” | inflammatory caf, icaf, chemokine-rich, msc-like |
| Fibro_myCAF | “mycaf”, “myofibroblast” | mycaf, myofibroblastic, acta2*, alpha-sma_ |
| Fibro_PVL | “pvl”, “perivascular”, “pericyte”, “smooth muscle” | pvl, perivascular, pericyte, mural, vsmc |
| Fibro_Cycling | “cycling”, “proliferating”, “mki67” | cycling caf, proliferating fibroblast |
| Endothelial | “endothelial”, “blood vessel” | endothelial, vascular, capillary |
| Mouse B cells (Lineage w = 0.5, State w = 0.5) | | |
| Mature_B | “mature_b”, “b cell” (prefix) | mature b, follicular b, b.fo, ms4a1* |
| Erythrocyte_like | “immature b” (mislabel), “erythrocyte” | erythrocyte, erythroid, rbc, hemoglobin-high |

| | | |
|---|---|---|
| Mast_like | “naive b” (mislabel), “mast” | mast cell, basophil-like, cpa3* |
| pDC_Myeloid_like | “precursor b” (mislabel), “pdc” | pdc, plasmacytoid dendritic, myeloid-like |

**Notes:**
* **Keyword triggers** denote **case-insensitive substring rules** applied to author-provided cluster labels to map them to harmonized GT categories (Methods).
† For the breast CAF/MSC benchmark, the dataset is prefiltered to stromal/vascular lineages; therefore, scoring emphasizes subtype resolution by setting **w_state = 0.7** and **w_lineage = 0.3**.

**Scoring.** We compute $S_{\text{anno}} = w_{\text{lineage}}\, s_{\text{lineage}} + w_{\text{state}}\, s_{\text{state}}$ with $s_{\text{state}} \in \{0,1\}$ and $s_{\text{lineage}} \in \{0,1\}$ by default. **Partial lineage credit ($s_{\text{lineage}} = 0.5$)** is enabled only for explicitly defined near-lineage pairs in the dataset-specific config (e.g., **Fibroblast ↔ Endothelial** in the CAF/MSC benchmark). Predictions in forbidden major-lineage sets (defined per benchmark) receive $S_{\text{anno}} = 0$. Full alias dictionaries are provided in **benchmarks/config**.

**Abbreviations. CAF**, cancer-associated fibroblast; **iCAF**, inflammatory CAF; **myCAF**, myofibroblastic CAF; **MSC**, mesenchymal stromal cell; **PVL**, perivascular-like; **vSMC**, vascular smooth muscle cell; **ISG**, interferon-stimulated genes; **IFN**, interferon; **MAIT**, mucosal-associated invariant T cell; **pDC**, plasmacytoid dendritic cell; **RBC**, red blood cell; **T_CM**, central memory T; **T_EM**, effector memory T; **T_EMRA**, effector memory RA; **T_EX**, exhausted T; **Tfh**, T follicular helper; **Treg**, regulatory T cell; **Th17**, T helper 17.

**Supplementary Table 4.** Conceptual comparison of LLM-based annotators and foundation models

| | LLM-scCurator (this study) | GPTCelltype[8] | CASSIA[9] | CellTypeAgent[10] | scGPT[11] |
|---|---|---|---|---|---|
| Intervention point | Pre-prompt marker distillation for annotation of user-defined clusters; optional hierarchical discovery (coarse-to-fine labeling) | Inference-time prompting (LLM wrapper) | Inference-time multi-agent reasoning (optional retrieval) | Inference-time candidates + expression-evidence verification | Model-centric foundation model (pretrain, fine-tune/mapping) |
| Requires user-supplied/precomputed marker table as the primary input? | No (does not require a precomputed marker table as the primary input) | Yes (expects precomputed markers/DEGs) | Yes (expects marker/DEG table) | Yes (expects markers + verification) | No (uses expression matrices; requires pretrained checkpoint + gene vocabulary) |
| Privacy / data exposure | Local-first; sends only compact cluster-level summaries (gene symbols + coarse context) | Sends marker summaries to the chosen LLM endpoint | Web UI or local run; exposure depends on deployment and endpoint | Local run + external evidence querying (as configured) | Local compute by default; optional hosted apps run on cloud GPUs (deployment-dependent). |
| Deployment / usability | pip; Docker/Apptainer; Colab; Seurat/Scanpy-compatible (export or reticulate workflows) | R/Seurat-friendly wrapper | Web UI + R/Python options | Python workflow (more moving parts) | PyTorch; GPU recommended at scale; browser-based cloud-GPU apps are available for mapping/annotation/ GRN. |
| Spatial applicability | Demonstrated on single-cell and spatial transcriptomics via cluster-level marker summaries | Typically used with scRNA-seq marker lists | Typically used with scRNA-seq marker tables | Typically used with scRNA-seq markers (with verification) | single-cell multi-omic tasks; spatial use depends on downstream integration |

| | | | | | |
|---|---|---|---|---|---|
| Unique advantage | Suppresses biological confounders and recovers identity markers via noise masking and Gini-informed distillation before prompting; outputs distilled marker lists usable by other LLM annotators; backend-agnostic (cloud or local). | Simple marker-to-label prompting | Trust via multi-agent critique / quality assessment | Trust via quantitative expression-based verification | Learns general representations via large-scale pretraining |

**Notes:**
Feature availability and data exposure depend on configuration (e.g., web UI vs local run; choice of LLM endpoint). Users should follow institutional policies for regulated data.

**Abbreviations: LLM**, large language model; **scRNA-seq**, single-cell RNA sequencing; **DEG**, differentially expressed gene; **GRN**, gene regulatory network; **GPU**, graphics processing unit.

**Supplementary Table 5.** Vendor pricing and approximate API costs for LLM-scCurator cluster-level annotation

| Backend model | Input price (USD/1M tokens) | Output price (USD/1M tokens) | Approx. cost/standard dataset (50 clusters), $ | Approx. cost/Atlas scale (200 clusters), $ |
|---|---|---|---|---|
| Efficiency | | | | |
| Gemini 2.5 Flash-Lite | 0.1 | 0.4 | 0.01 | 0.04 |
| GPT-5 Mini | 0.25 | 2 | 0.06 | 0.24 |
| Gemini 2.5 Flash | 0.3 | 2.5 | 0.07 | 0.28 |
| Performance | | | | |
| Gemini 2.5 Pro | 1.25 | 10 | 0.28 | 1.12 |
| Gemini 3 Pro (preview) | 2 | 12 | 0.35 | 1.4 |
| GPT-5.2 | 1.75 | 14 | 0.39 | 1.56 |

**Notes**:
Costs are estimated assuming a conservative budget of 1,000 total tokens per cluster query (≈500 input + 500 output). This represents an upper bound to accommodate extended reasoning; actual token usage for concise JSON outputs is typically lower.
Prices reflect vendor rates as of December 2025 and assume standard datasets (~50 clusters) or atlas-scale runs (~200 clusters), respectively.

**Supplementary References**

1. Tirosh, I. *et al.* Dissecting the multicellular ecosystem of metastatic melanoma by single-cell RNA-seq. *Science* 352, 189–196 (2016).
2. Zheng, L. *et al.* Pan-cancer single-cell landscape of tumor-infiltrating T cells. *Science* 374, abe6474 (2021).
3. Wu, S. Z. *et al.* A single-cell and spatially resolved atlas of human breast cancers. *Nat. Genet.* 53, 1334–1347 (2021).
4. Tabula Muris Consortium. A single-cell transcriptomic atlas characterizes ageing tissues in the mouse. *Nature* 583, 590–595 (2020).
5. Furudate, K. *et al.* Spatial colocalization and molecular crosstalk of myofibroblastic CAFs and tumor cells shape lymph node metastasis in oral squamous cell carcinoma. *PLoS Genet.* 21, e1011791 (2025).
6. Oliveira, M. F. de *et al.* High-definition spatial transcriptomic profiling of immune cell populations in colorectal cancer. *Nat. Genet.* 57, 1512–1523 (2025).
7. Crowell, H. L. *et al.* Orchestrating spatial transcriptomics analysis with Bioconductor. *bioRxivorg* (2025) doi:10.1101/2025.11.20.688607.
8. Hou, W. & Ji, Z. Assessing GPT-4 for cell type annotation in single-cell RNA-seq analysis. *Nat. Methods* 21, 1462–1465 (2024).
9. Xie, E. *et al.* CASSIA: a multi-agent large language model for automated and interpretable cell annotation. *Nat. Commun.* (2025) doi:10.1038/s41467-025-67084-x.
10. Chen, J., Zhang, J., Yao, H. & Li, Y. CellTypeAgent: Trustworthy cell type annotation with Large Language Models. arXiv:2505.08844, (2025).

11. Cui, H. *et al.* scGPT: toward building a foundation model for single-cell multi-omics using generative AI. *Nat. Methods* 21, 1470–1480 (2024).
